# Finite element model of the non-keratinized buccal tissue under the impact of negative pressure

**DOI:** 10.1101/2024.11.14.623564

**Authors:** David Klein Cerrejon, Daniel Gao, David Sachs

**Affiliations:** Institute of Pharmaceutical Sciences, Department of Chemistry and Applied Biosciences, ETH Zurich; 8093 Zurich, Switzerland; Citus AG, Ueberlandstrasse 129, 8600 Dübendorf, Switzerland

## Abstract

The buccal mucosa is a highly interesting site for non-invasive drug delivery due to its relatively permeable epithelium and good accessibility. Recently, device-based systems have enabled the delivery of macromolecular drugs by leveraging mechanical stretching forces on the tissue to assist drug diffusion. Despite the successful exploitation of the buccal route with such systems, the biomechanics of buccal tissue are still poorly characterized and understood due to a lack of adequate characterization methods. Therefore, we propose a combination of physiological tissue modeling with simple suction experiments as a tool for characterizing and understanding the buccal tissue under the impact of negative pressure. Here, we present an initial step towards a multiphasic and poroelastic model specifically designed for the non-keratinized buccal tissue under the impact of negative pressure. A validated finite element model (FEM) for human skin was adapted to represent the histological structure of porcine buccal tissue. We performed suction experiments using the NIMBLE device, specifically developed for measuring skin stiffness, to characterize its mechanical behavior and train the FEM model. The resulting simulation tracks essential physiological parameters and allows the prediction of measurable changes in the tissue, such as the thinning of the epithelium and single-cell stretching. The FEM simulation was validated through histochemically stained tissue sections at the NIMBLE application site. A good correlation was demonstrated between predicted and experimentally observed changes. This work serves as a first step towards a computational representation of buccal tissue under the impact of negative pressure.

## Introduction

The buccal mucosa is a highly interesting site for noninvasive drug delivery due to its ease of accessibility, relatively good permeability, mild conditions (i.e., pH and low proteolytic enzyme activity), and the avoidance of the hepatic first-pass metabolism (*1–4*). However, the buccal route is currently restricted to the administration of low molecular weight drugs due to the lack of permeability of macromolecules (*3*). Drug delivery systems leveraging physically assisted systems such as negative pressure (*4*), liquid jets (*5*), needle injectors (*6*), iontophoresis (*7*) and focused ultrasound (*8*) in combination with advanced drug formulations have overcome some of these limitations, resulting in more effective delivery of small and macromolecular drugs through the buccal mucosa (*9*). Although these systems are heavily dependent on the mechanical response of the mucosa towards mechanical forces such as stretching, the biomechanics of buccal tissue and response to mechanical triggers remain inadequately explored due to a lack of suitable characterization methods. Generally, the buccal tissue is organized in a layered structure, starting with the non-keratinized epithelium that represents the key diffusion barrier and comprises around 50 layers of tightly packed stratified squamous cells (*1*, *10–12*). The epithelium and lamina propria, the layer beneath, are separated by the basal layer and membrane – a robust assembly of basal cells and extracellular matrix proteins serving as an anchor for the lamina propria. The lamina propria encompasses blood vessels, nerves, fibroblasts, and mast cells embedded in a dense collagen network (*11*, *13*). This network transitions to the submucosa, consisting of fibrocollagenous and elastic tissues, followed by the muscle layer (*11*, *13*). Buccal tissue shows high deformability with stiffening occurring at large strains in both uniaxial tensile tests and compression tests (*14*). Dynamical mechanical tests revealed dissipative and time-dependent behavior, which is explained by the dissipative behavior of the collagen network and interstitial fluid flow (*15*, *16*). However, the measured mechanical properties of the mucosal tissue strongly vary depending on the method, sample type and preparation. For example, determined Young’s moduli range from a few kPa in atomic force microscopy (AFM) to several MPa in uniaxial tension tests (*17*, *18*). Mechanical properties may differ between different layers due to varying cell types and morphologies (*13*). Thus, measurement outcomes may strongly change when measuring tissue samples of different thicknesses or using methods on isolated individual layers (*19*). Also, currently used stiffness measurement techniques, such as AFM or tensile testing, are invasive or impractical and cannot be carried out on a living human (*17*). Therefore, the physical characterization of the buccal tissue is currently restricted to *ex vivo* tissue studies (*17*). Despite these limitations and uncertainties in the tissue characteristics, hyperelastic, non-linear models with dissipative and biphasic features have been developed to investigate the mechanical properties of the buccal tissue (*18*, *20*, *21*) As a consequence of limited input data and the absence of validation tools, their capacity to represent the complex nature of buccal tissue is constrained. Moreover, the contribution of individual tissue layers, stemming from different mechanical properties and fluid movement across different layers, is not respected. Novel approaches such as the NIMBLE try to overcome these limitations by utilizing noninvasive and simple negative pressure applications, allowing for measurements in living humans (*22*). The combination of different experimental protocols, including different NIMBLE probe dimensions, allows for the characterization of individual tissue layers (*22*). The integration of these combined methods has yielded an improved characterization of both skin and cervical tissues and is currently being translated for clinical applications (*19*, *22–25*). Incorporating NIMBLE measurements and insights from the FEM enables the translation of a skin model into an initial *in silico* model for buccal tissue, allowing simulation of its response to negative pressure. This approach may be combined with drug delivery technologies, such as a suction patch, to analyze the effects of negative pressure on tissue during application. (*4*).

This work proposes a multilayer mechanical model of buccal tissue. Therefore, we adapted a validated FEM of the skin (Figure 1A) to the histology of the buccal tissue. Second, we performed suction experiments with the NIMBLE device to characterize its mechanical behavior (Figure 1B), and adapted the multiphasic material model, differentiating mechanical parameters in the epithelium, lamina propria, and muscle (Figure 1C). Third, we experimentally quantified epithelium as well as cell deformation and compared them to the modeling predictions of porcine buccal tissue. (Figure 1D). Finally, we investigated the behavior of the proposed material model and the biomechanical implications of negative pressure on the tissue (Figure 1D).

**Figure 1.**
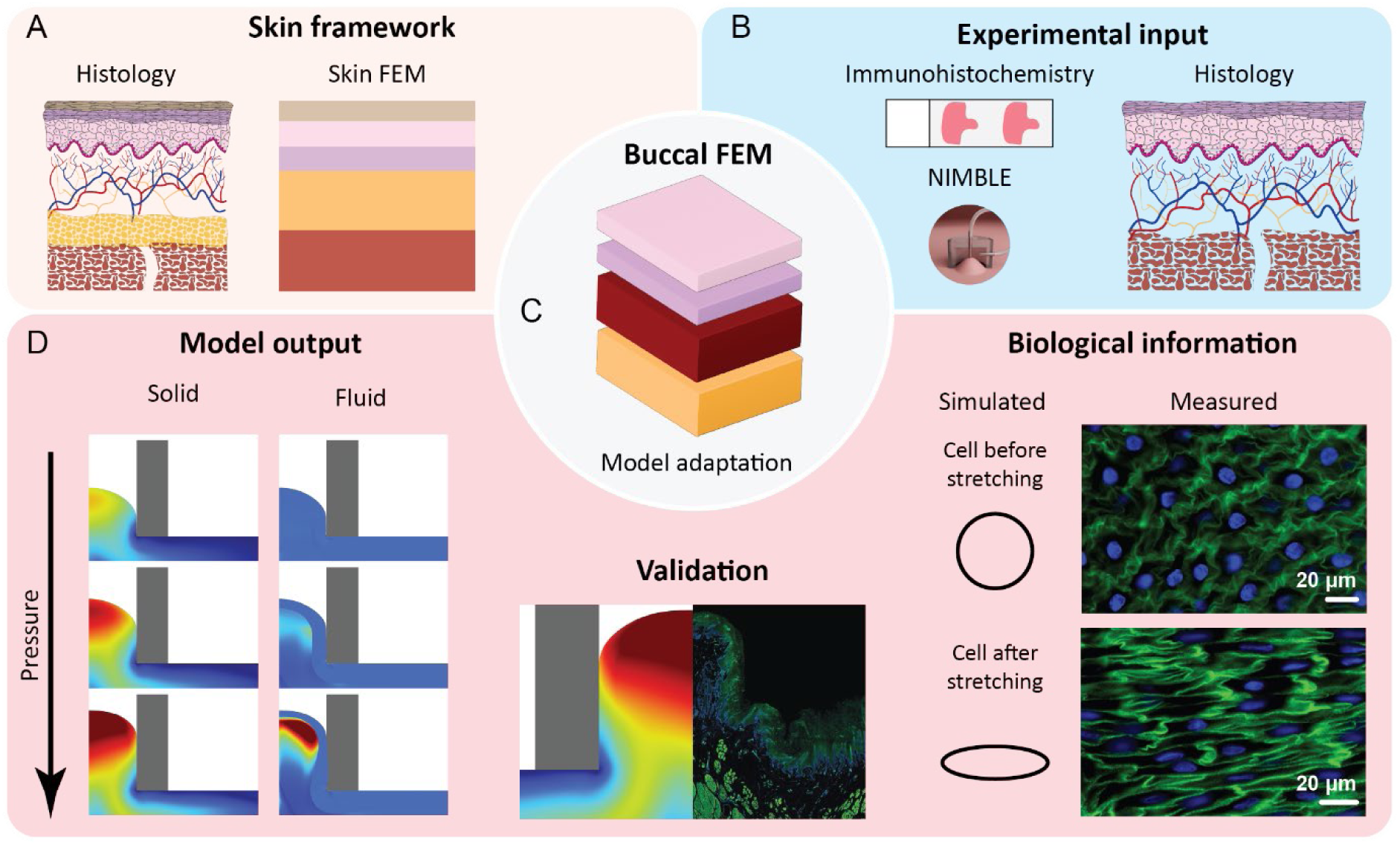
Graphical abstract of the development and validation of a buccal tissue FEM. A) A skin FEM was adapted according to B) the histology and experimental input data to obtain C) a new buccal tissue FEM. D) The FEM output was evaluated and compared to experimental data of an independent set of experiments.

## Results and discussion

### Suction measurements on buccal tissue

First, the parameters of the multiphasic model were fitted with the experimental results of deformation height measurements with the NIMBLE device (Figure 2, inset). Using the derived model parameters (Table S1), the simulation successfully replicated the experimental suction measurements. The buccal mucosa exhibited a very soft behavior with a deformation of 3 mm occurring at a negative pressure of 1.8 ± 1.1 kPa. The required negative pressure increases non-linearly for higher elevations, reaching approximately 7.8 ± 4.4 kPa for 4 mm and 21.5 ± 4.5 kPa for 5 mm. The stiffness of individual tissue layers was determined by an inverse analysis. The initial solid volume fraction (*26*) and permeability of interstitial fluid in the lamina propria (*27*, *28*) were selected based on literature findings. Additionally, osmotic pressure was incorporated to account for the presence of glycosaminoglycans and proteoglycans in the same order of magnitude as observed in the dermis (*26*, *29*, *30*). The muscle layer was set proportional to the lamina propria and chosen to be one order of magnitude stiffer than the lamina propria. The remaining parameters were fitted iteratively to match the experimental response. The final parameters were listed in Table S1. The obtained material characteristics fit the experimental data for 3, 4, and 5 mm elevations. Interestingly, between 4 and 5 mm deformation, the FEM predicted a jump in the pressure deformation curve. This may be explained by activation of the drastically stiffer muscle tissue, where a negative pressure of 3.4 ± 0.2 kPa (n = 3 tissue samples) was required to reach a deformation of 2 mm or the non-linearity. Young’s moduli of the epithelium, lamina propria, and muscle were obtained by computational uniaxial tests with the experimentally fitted FEM (Table S2).

**Figure 2.**
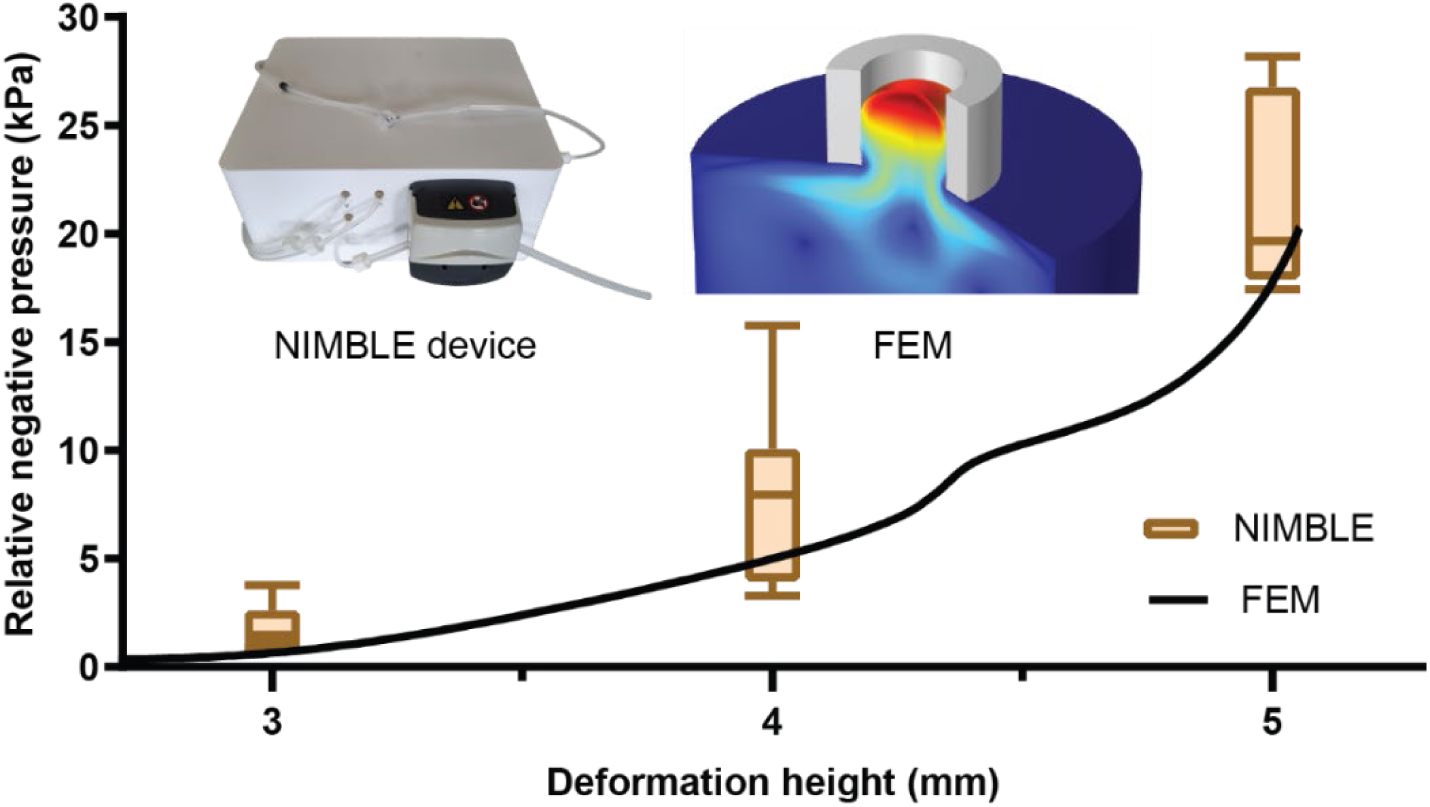
The deformation height as a function of relative negative pressure for experimental measurements and refined FEM simulation. The black line indicates the output of the optimized FEM. The box-whisker-plot (box: 25 – 75 percentile, whisker: min – max) shows the pressure needed at a defined NIMBLE probe height (n = 6 tissue samples with triplicate measurements at two different positions). The inserted graphic depicts the NIMBLE device and an exemplary 3D view of the solid deformation in simulation with a deformation height of 5 mm. The color gradient (blue to red) indicates the degree of solid deformation from its original position.

In the uniaxial computation, a cuboidal sample was elongated quasistatically up to a strain of 0.5 (Figure 3). The non-linearity of the Rubin-Bodner model was well captured, especially for the lamina propria and muscle (Figure 3 A – B). Therefore, the tangent modulus for low (Figure 3C, left) and high (Figure 3C, right) strains were several orders of magnitude apart, a well-known behavior of soft biological tissue. The model predicts Young’s moduli for small strains (1 – 10%) in the order of 10^-2^ kPa, while for high strains (40 – 50%) values can reach several kPa up to MPa for the muscle layer. Literature values range from less than one kPa to several MPa, depending on analysis and tissue preparation method (*17*, *18*). Compared to other biological tissues, buccal tissue exhibited a much softer behavior, with a Young’s modulus more than three orders of magnitude lower than that of skin. (*23*, *31–33*). Our analysis revealed that, while the reported values for large strains fell within the expected range (kPa to MPa), the values for small strains were lower. Accurately predicting Young’s moduli at low strain is challenging due to difficulties in defining the initial configuration, specifically the forces and stretches at the beginning of the experiment. This challenge is particularly relevant for very soft tissues, where slight variations in threshold force can result in large deformations and substantial changes in the initial configuration. Since no threshold force was applied in the suction experiments, the initial conditions of this inverse analysis may differ from those reported in uniaxial experiments. Secondly, deformations in suction experiments can also occur due to motion (slipping) of the tissue, rather than stretching, which would lead to an underestimation of the stiffness for small stretches compared to uniaxial tests. Moreover, tissue preparation with regards to tissue thickness, viability, and post-mortem stiffening can affect the measured stiffness.

**Figure 3.**
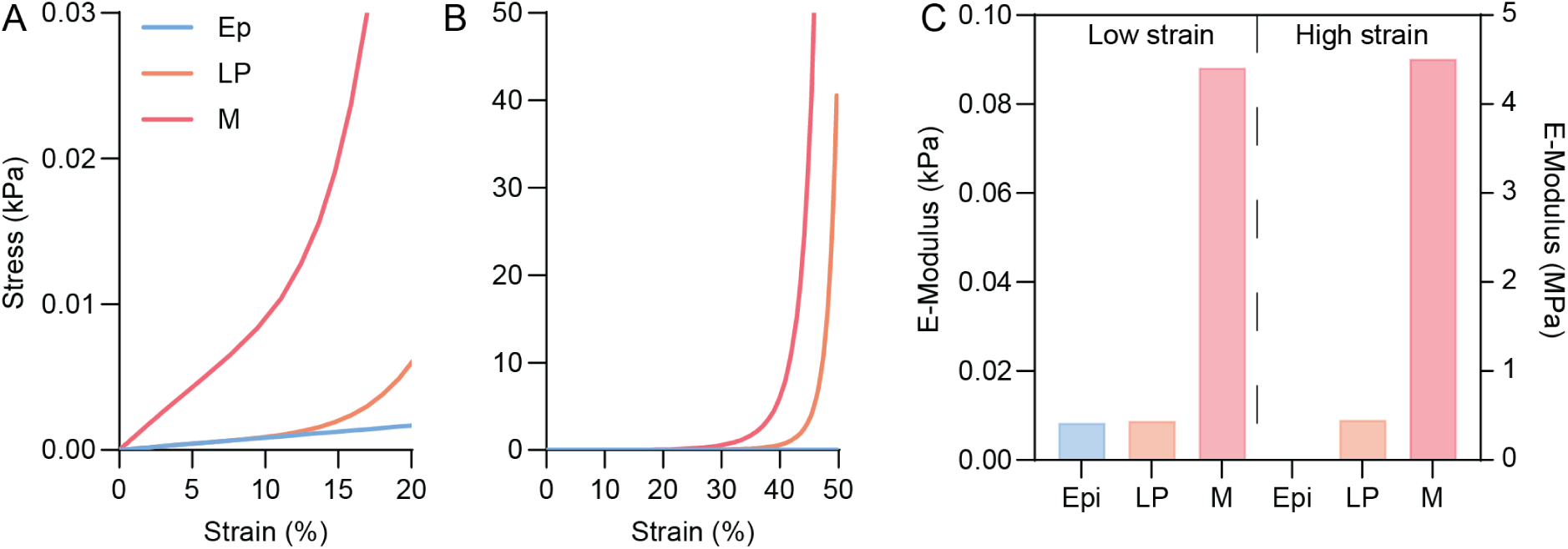
Computational uniaxial test of individual layers in the fitted FEM. Stress-strain curves for A) low strain and B) high strain. C) Young’s modulus for low strain in the range of 1 – 10% and high strain 40 – 50%. Abbreviations: Ep: epithelium (blue); LP: lamina propria (orange); M: muscle layer (red).

### Experimental validation and cell deformation

A separate set of experiments was performed in order to assess the model’s capability to predict local tissue response. The epithelial thinning and deformations of single cells in the epithelium were characterized experimentally and compared to the predicted values obtained by the trained model. Hence, NIMBLE probes of varying heights (3 – 5 mm) were applied to the buccal mucosa until maximal deformation was reached. The tissue, along with the NMBLE probe, was then snap-frozen to preserve the induced deformation. Subsequent processing and histochemical staining with iFluor^TM^488-conjugated phalloidin (green) for cytoskeleton labeling (binding to F-actin) and Hoechst 33342 (blue) for nuclear staining (Figure 4A - C) was performed. The basal layer, characterized by densely aligned blue-stained cell nuclei, clearly separates the epithelium from the lamina propria. Underneath the lamina propria, only little F-actin containing tissue components were observed, which at the lower end gradually transitioned to the muscle, rich in F-actin. The clear separation of the epithelium enabled the measurement of its thickness within (center) and outside (outside) of the deformation. Outside, the epithelium exhibited an almost unstretched state, representing the native tissue condition. The average epithelial thickness was 1.3 ± 0.4 mm, and the shape of individual cells within the epithelial layer appeared rather circular/elliptical (Figure S1B and D). In contrast, at the center of the NIMBLE probe deformation, the tissue underwent significant stretching. The epithelial thickness was obtained by applying the Cauchy-Green tensor, extracted from the center 0.3 mm below the surface, to an epithelial thickness of 1.5 mm. It decreased with increasing probe height and negative pressure with an average deviation of 17% to the experimental mean, which was close to the experimental error (average SD of 14%) (Figure 4G). The deformation behavior of the lamina propria and the muscle under negative pressure-induced deformation visually matched the simulated deformation of both layers (Figure 4A – F). Furthermore, the cellular shape was quantitatively compared in both regions of interest (center and outside of the application side). The average height over the length of single cells (n = 10) was compared (Figure 4H). The center line indicates a perfect circular shape. Non-stretched epithelial cells (grey circle) exhibited a more circular appearance, while those at the center of the deformation displayed significant elongation (blue triangle) (Figure S1C and D). The degree of elongation in the x-direction (X) of cells increased with the height of the NIMBLE probe by up to 226% of its initial value and contracted to 37% of their initial height (y-direction (Y)) (Figure 4H).

**Figure 4.**
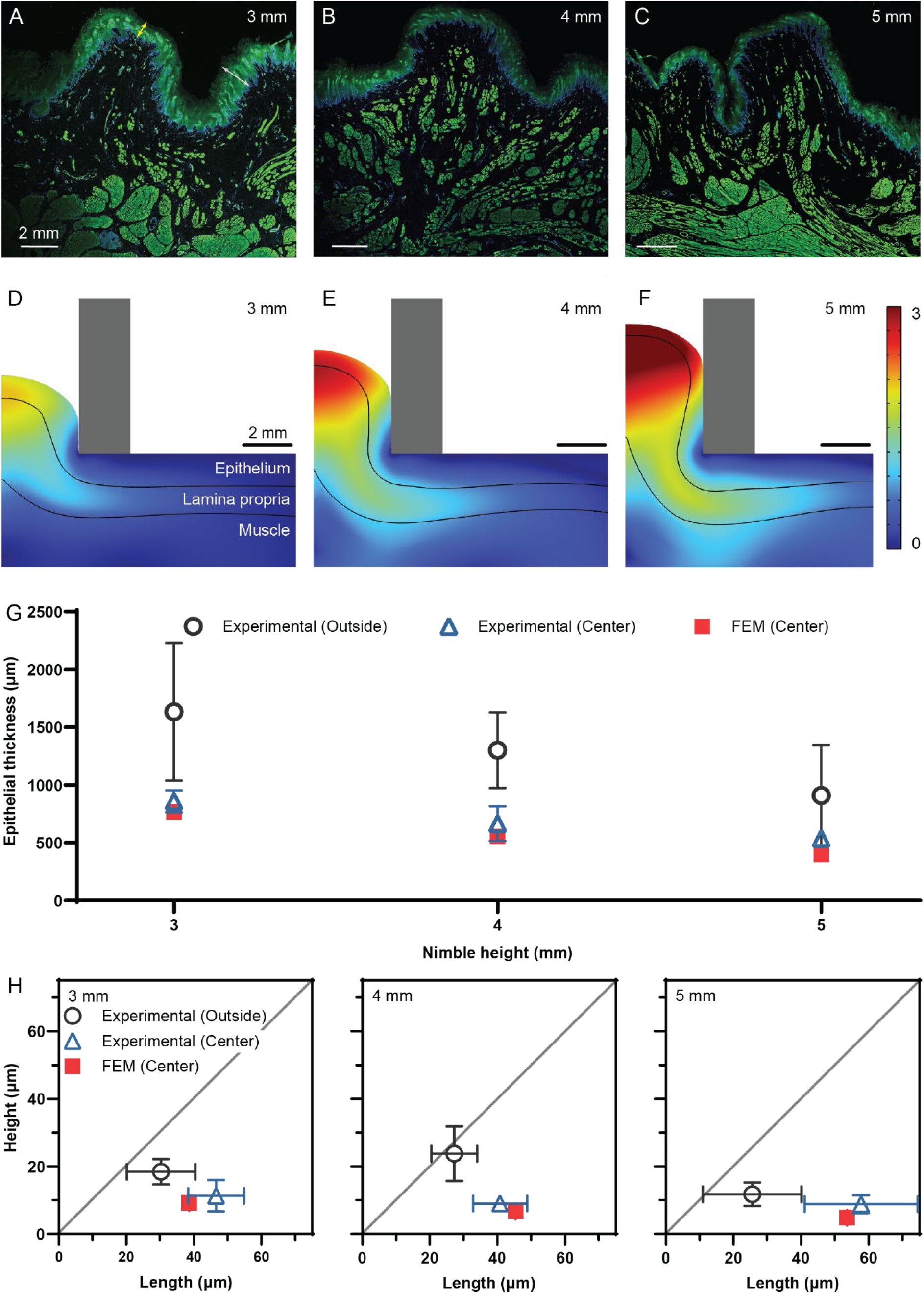
*Ex vivo* validation of the FEM with histochemically stained porcine tissue samples. Histochemically stained slides of snap-frozen porcine buccal tissue during NIMBLE application after reaching closing pressure (A – C). Three different heights of the NIMBLE probe (i.e., A) 3 mm, B) 4 mm, and C) 5 mm) were compared. To identify different tissue layers and the outline of single cells, the cytoskeleton (F-actin, green) and nucleus (blue) were stained. The FEM simulated solid deformation for individual layers for a height of D) 3 mm, E) 4 mm, and F) 5 mm. The color gradient indicates the deformation of the tissue from its initial configuration and the solid black lines indicate the separation of individual tissue layers. G) Measured thickness of the epithelial outside (grey, circle) and in the center (blue, triangle) of the NIMBLE application side at different deformation heights (n = 5 measurements, error bar indicate SD), compared to the thickness obtained by the FEM in the center (red, square), when the predicted Cauchy-Green tensor (vertical direction) was applied to a thickness of 1.5 mm. H) Cell dimensions (length and height) measured (n = 10 cells, error bars indicate SD in length and height) outside (grey, circle) and in the center (blue, triangle) of the NIMBLE application side compared to the dimensions (red, square) obtained when the stretching coefficient from the FEM was applied to a rather circular shape with the average initial dimension of the cell (X = 27.7 µm and Y = 17.9 µm) (red, square). The solid diagonal line indicates an ideal circle.

In the FEM, the cell stretching was obtained by applying the Cauchy-Green tensor to an average cell that was defined as an ellipse described by length (X = 27.7 µm) and height (Y = 17.9 µm) (experimentally measured, n = 3 samples and 10 cells/sample). Similar to the experimental data, the calculation resulted in an elongation in X and compression in Y (Figure 4H). The FEM predicted the cell deformation with a deviation of < 26% (X and Y) for 3 and 4 mm elevation compared to the average experimental value. At the elevation of 5 mm, it showed a similarly low deviation in X (7%) and overestimated the compression to the final height of 4.7 µm instead of the experimentally determined 8.7 ± 2.7 µm (45% deviation). Differences between numerical simulation and experimental results show that while an incompressible single-phase material is a good first approximation to describe the deformation of cells, further studies are required to refine the model. These should, on the one hand, investigate the material model chosen for the cellular layer and, on the other hand, validate the incompressibility, degree of slipping, and single-phase assumptions.

### The mechanical behavior of buccal tissue

The FE model allowed for analysis of the mechanical behavior of individual layers and their interaction. Figure S2 depicts circumferential (A), radial strains (B), and the volume change (C) of a cross section of the tissue during a suction experiment at an apex displacement of 4 mm. Biological tissues modeled with a biphasic approach show volume increase due to volume influx, or volume decrease due to fluid efflux (*30*). The NIMBLE probe was presented as a cross section in grey. The different modeling approaches i.e. incompressible (epithelium) *vs.* compressible (lamina propria and muscle) resulted in distinct behavior during the suction experiment. The epithelium showed large positive strains in the radial direction (1.5 – 3: Up to four times the initial value) and compressive strains (-0.5: half the initial value) in the circumferential direction (Figure S2A and B). In contrast, the lamina propria showed positive strains in both radial (Figure 7A) and circumferential direction (Figure S2B), resulting in an overall volume increase of up to 2.6 times (Figure S2C). Simulated volume changes of the lamina propria were in line with the swelling observed *ex vivo* (Figure 4A – D). The multiphasic and multilayered approach allowed to differentiate the response of individual layers to the negative pressure induced stretching. Although qualitative trends align with the experimental data, the absolute values, especially at the interface of tissue layers, must be evaluated with caution, as no further quantitative experimental validation was carried out in this study. To the best of our knowledge, no comparable literature is available.

### Suction induced flow and change of permeability in the lamina propria and muscle

To better understand the buccal tissue response to long term negative pressure we simulated the NIMBLE application for 10 min and then released the pressure. We tracked the pressure, displacement, volume changes, the magnitude of interstitial fluid flow velocity, and deformation induced permeability changes (Figure 5A – D) over 1500 s in the lamina propria. The changes in permeability are normalized with respect to the initial permeability. In the simulation, the tissue was initially elevated by a negative pressure of 0.6 kPa within 2 s and held at 0.6 kPa for 600 s. The spatial-temporal evolution of volume change, interstitial fluid flow velocity, and permeability changes (t = 2, 5, 10, 150, and 300 s) were plotted in Figure 5E – G. As the epithelium was assumed to be an incompressible solid, no fluid flow, volume change or change in permeability of the interstitial fluid was reported for the epithelium (purple).

**Figure 5.**
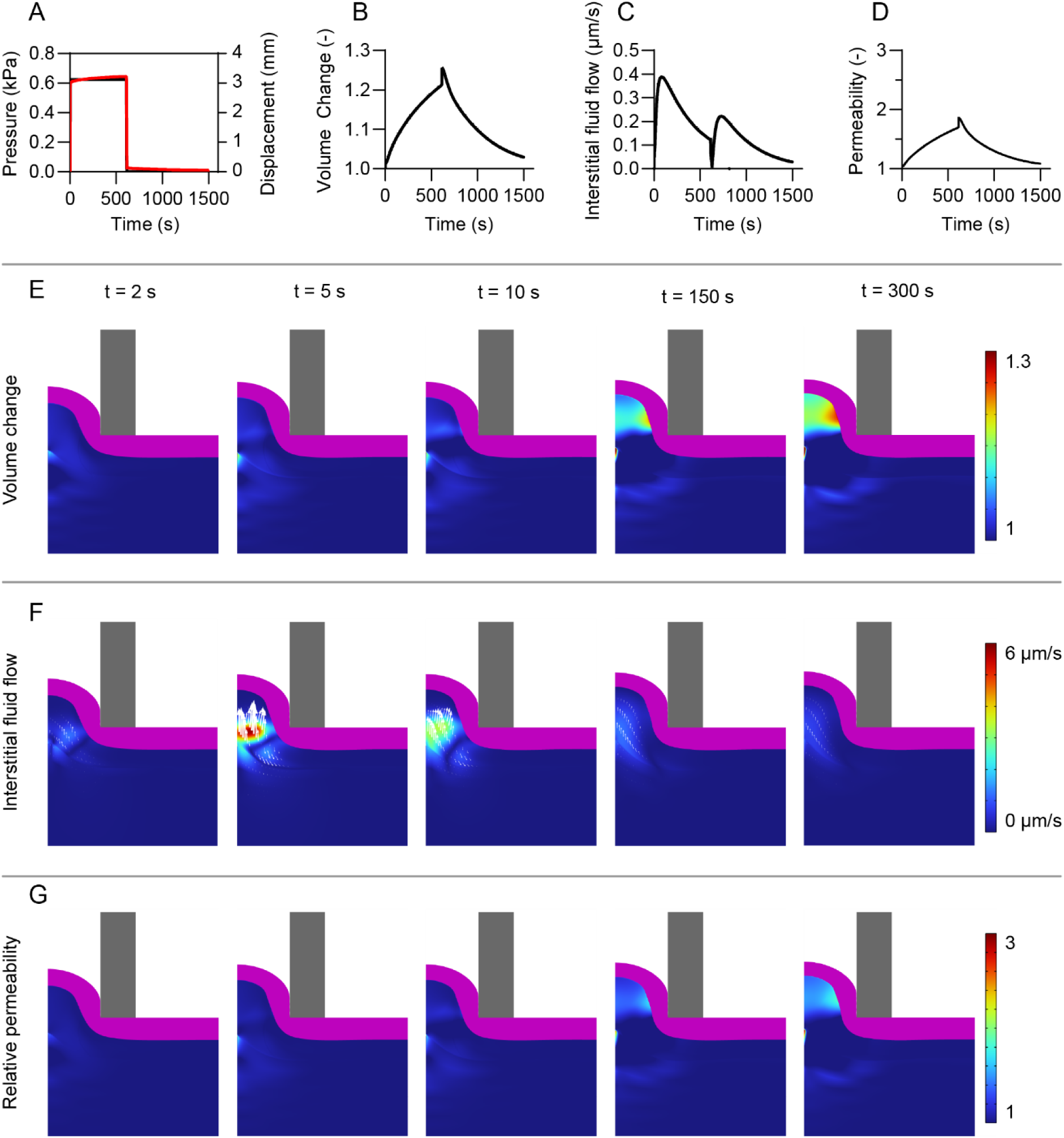
Tissue behavior predicted by the finite element simulation after a prolonged application of a negative pressure. A) A negative pressure ramp (black) up to 0.6 kPa was reached after 2 s and applied for 10 min before being released within 4.9 s. The corresponding apex displacement is shown in red. B) Volume change in the center of the tissue at the boundary of epithelium and lamina propria. C) Magnitude of interstitial fluid flow velocity at the same position as in B. D) Change in permeability at the same position as in B. The spatial-temporal response is depicted by a cross section at 2, 5, 10, 150, and 300 s for the change in E) volume, F) interstitial fluid flow and direction and G) relative permeability. Please note that in E) and G) at the interface of lamina propria and the muscle at the center a minor computational artefact is present. The thickness of the grey NIMBLE probe indicates 2 mm.

Although the full deformation of 3 mm was achieved immediately, volume changes continued to increase up to 1.3 times compared to its initial size. The change in volume was rapid within the first 10 s, but the rate of change decreased over time (Figure 5B, E). This can be explained by the interstitial fluid flow towards the apex, which peaked with a slight offset around 5 s (Figure 5F) and continued at a reduced flow rate, resulting in a swelling of the lamina propria over time. The relative permeability increased with application time to up to 3 times of its initial value (Figure 5G). The largest volume changes were predicted and experimentally observed in the lamina propria (Figure 4A and 5E). The lamina propria represents the softest region of buccal tissue, consisting of a comparably loose collagen network and allowing for comparably good interstitial fluid flow (*11*). This allows for higher changes in volume and increase in permeability, compared to drastically stiffer and denser muscle. After 600 s, the pressure was reduced within 4.9 s to 0 kPa and held at 0 kPa thereafter. When the pressure was released, the apex displacement returned immediately close to zero (> 95% of max. displacement) (Figure 5A and 6). The volume and permeability decreased simultaneously with the negative pressure. The outflux of interstitial fluid reached a maximal flow rate of 3 µm/s (Figure 6B) but again continued for more than 5 min, before completely vanishing. The simulation appeared to reflect the natural behavior well, as a similar prolonged recovery time from a negative pressure impact was observed with the application of a 6 mm height suction cup on human buccal mucosa for 30 min (*4*). After the suction patch removal by the volunteers, the buccal tissue relaxed almost entirely to its original position, but some swelling at the application site remained before completely vanishing after 1 h (*4*). This may be attributed to a combination of viscoelastic deformation of the solid and the prolonged outflow of interstitial fluid from the application side.

**Figure 6.**
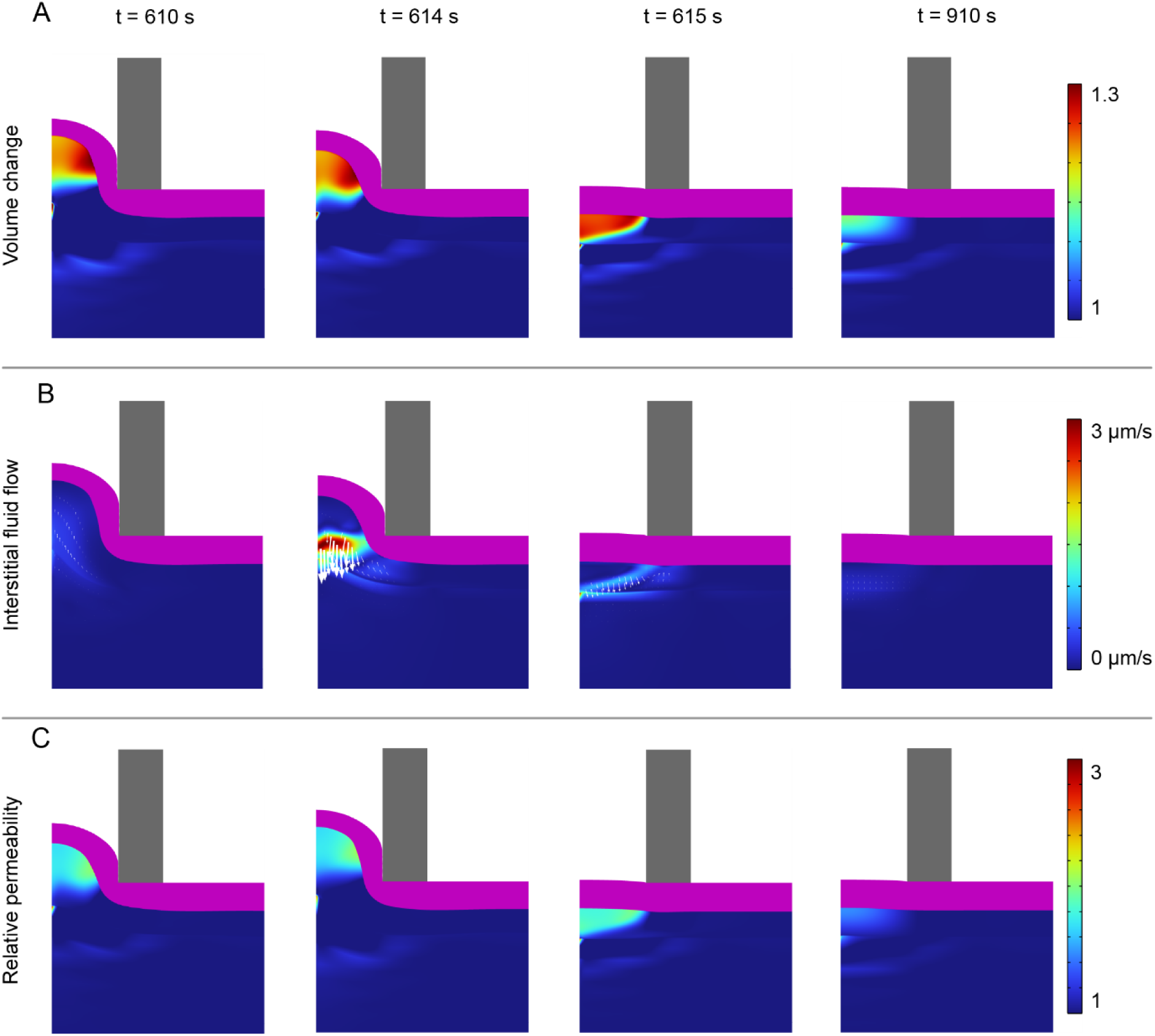
Tissue behavior predicted by the finite element simulation after a prolonged application of negative pressure. To evaluate the spatial-temporal response, a cross-section of the tissue is depicted at 610, 614, 615, and 910 s for the change in A) volume, B) interstitial fluid flow and direction, and C) relative permeability. Please note that in A) and C), at the interface of lamina propria and the muscle at the center, a minor computational artifact is present. The thickness of the grey NIMBLE probe is set at 2 mm.

## Conclusion

We report the development and proof of concept for a multiphasic and poroelastic buccal tissue FEM. The previously reported and thoroughly validated skin FEM was adapted to account for the mechanical and histological differences of the buccal tissue. Fresh porcine buccal tissue was used to evaluate mechanical parameters and train the FEM by using the NIMBLE device. It was found that buccal tissue was at least three orders of magnitude softer than skin. With an independent validation data set, we analyzed the epithelial stretching caused by negative pressure using the NIMBLE device and assessed the stretching of individual cells in the epithelium through histochemical staining of tissue slides. We observed the thinning of the epithelium and a longitudinal elongation of up to > 200% and transversal compression to < 40% of an individual cell depending on the degree of stretching. The FEM predicted qualitative changes in individual tissue layers and suggested a physiological behavior that is in line with the previously observed impact of buccal suction patches. However, further validation, larger sample size, in-human experiments and measurement of physiological parameters still need to be obtained to predict the tissue behavior quantitatively and reliably in humans. This work serves as a first step towards a computational representation of buccal tissue under the impact of negative pressure. Moreover, in combination with devices utilizing the buccal mucosa such as the suction patch, it has the potential to be used as a digital twin to better understand the impact on the tissue and guide the development of new technologies.

## Material & Methods

Phosphate-buffered saline (PBS), bovine serum albumin ≥ 96% (BSA) (≥96%) Triton X- 100, methanol-free paraformaldehyde (PFA) and 2-methylbutane were purchased from Sigma-Aldrich. Dulbecco’s Modified Eagle Medium with F-12 Nutrition Mixture (with glutamine and 15 mM HEPES, without phenol red) (DMEM/F12) and antibiotic-antimycotic (100×) (10,000 U of penicillin, 10 mg of streptomycin, and 25 μg of amphotericin B per mL) were purchased from Thermo Fisher Scientific. Hoechst 33342 dye and ProLong Diamond Antifade Mountant was purchased from Invitrogen. iFluor 488–conjugated phalloidin (ab176753) was purchased from Abcam. Optimal cutting medium (OCT) was purchased from Leica Microsystems.

### Tissue preparation for *ex vivo* analysis

Porcine buccal tissue (entire cheek collected) (animal age: 5 – 6 months, weight: 100 – 120 kg) was obtained from the slaughterhouse (Schlachtbetrieb Zürich AG). The tissue was immersed in DMEM/F12 containing 1% (v/v) antibiotic-antimycotic 100X immediately after extraction and placed on ice within 30 min. The cheek was used no later than 3 h after collection.

### Suction measurement

*Ex vivo* mechanical tests were carried out on 6 porcine buccal tissue samples from different animals. The entire cheek was prewarmed in DMEM/F12 containing 1% (v/v) antibiotic-antimycotic 100X to 37 °C before use. For the characterization of the mechanical behavior, suction experiments with the NIMBLE were performed (*22*). Probes with probe opening diameters of 6 mm and pin heights of 3, 4, and 5 mm were used. Each specimen was measured at two locations in triplicates (Figure S3A). When the suction deformation reached the pin height, a pressure difference could be detected between the probe and reference sensor. At 1 kPa difference, the pressure of the probe was extracted. The system could measure a maximal negative pressure of 60 kPa.

### Multiphasic model of buccal tissue

Due to the layered structure of the buccal tissue, we distinguished several layers in the tissue model by applying different hyperelastic strain energy function parameters (Figure S3B and C). All layers were modeled in the framework of nonlinear elasticity. The deformation gradient *F* (Eq. 1) results from the mapping *χ* which maps point from its initial configuration *X*(*t*) to its current configuration *x*(*t*).

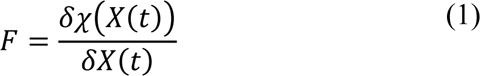

#### Epithelium

The cellular epithelial layers were modeled as a single phase incompressible neo-Hookean material. The strain energy density per reference volume is given by Eq. 2:

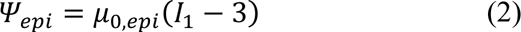

Therein, μ_0,_*_epi_* represents a material parameter and *I*_1_ the first invariant of the deformation tensor.

The epithelium was treated as incompressible as it consists of densely packed cells that were assumed as water (incompressible) filled shells that preserve their volume even under stretching (*11*, *34*, *35*). However, to the best of our knowledge this has never been investigated for non-keratinized mucosal epithelium and not for cells undergoing large strain as experienced in the presented experiments. Therefore, further investigations need to be conducted in order to confirm this assumption.

#### Lamina Propria and Muscle

The lamina propria and the muscle were modeled as a biphasic material consisting of solid and liquid phases. Assuming incompressibility of both phases, the mixture behaves compressible by fluid efflux or influx through the boundaries. The contribution to the strain energy from the solid component was modeled using a Rubin-Bodner strain energy density (*36*) (Eq. 3):

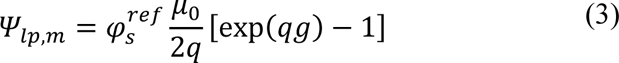

The function *g* is thereby given as (Eq. 4)

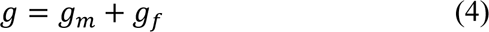

With *g_m_* and *g_f_* corresponding to the nonlinear elastic contributions of the matrix and the fibers. The specific functions read (Eq. 5a, 5b):

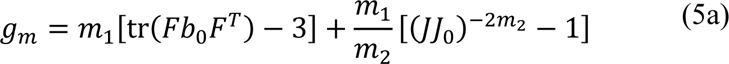

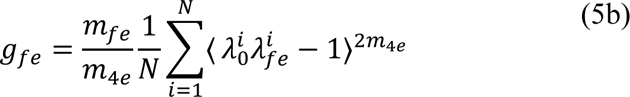

The parameters *m*_1_, *m*_2_, *m_fe_* and *m*_4*e*_ represent material parameters. In the strain energy function, the first term is similar to a Neo-Hookean solid, which was represented by the Cauchy deformation tensor *b* = *FF^T^* and the second term represents energy contributions due to changes in volume, represented by the third invariant of the deformation tensor *J* = *det* (*F*). The fibrous component was added to account for possible anisotropy in the material due to collagen and elastin fibers. In this specific case, we modeled *N* = 32 fiber families. The Macaulay brackets ⟨⋯ ⟩ ensure that fibers were only active in tension.

The motion of the interstitial fluid was governed by the fluid chemical potential *μ_f_*, which was given by (Eq. 6):

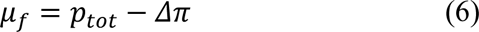

The fluid chemical potential was the difference of the hydrostatic pressure *p_tot_* and the osmotic pressure *Δπ*. The interstitial fluid velocity *q* followed Darcy’s law (Eq. 7):

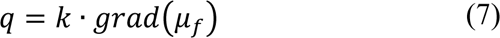

The proportionality tensor *k* = *k*_0_*I* described the permeability tensor of the interstitial fluid and was assumed to be isotropic in the current configuration. The governing equations for the biphasic solid were given by the conservation of mass and the conservation of momentum. The conservation of mass (Eq. 8) with the spatial velocity of the solid *v_s_*:

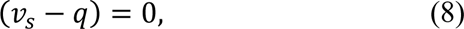

The conservation of momentum was given by (Eq. 9):

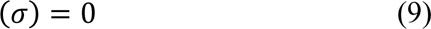

The Cauchy stress tensor *σ* results from the sum of the solid contribution to the strain energy function and the hydrostatic pressure (*23*).

### Finite element implementation

The model was implemented into the commercial finite element software COMSOL Multiphysics (COMSOL 6.0, COMSOL Multiphysics). The NIMBLE was modeled using a linear elastic material, with a large Young’s modulus (Young’s modulus = 210 GPa). An axisymmetric model was chosen to reduce computational effort, exploiting the symmetry of the system. The penalty factor method was used to simulate the contact condition, with the penalty parameter varying from 20 to 200. A negative pressure boundary condition was applied on the epithelium. The bottom side of the muscle was fixed, while all other boundaries were free to move. No fluid flow was allowed from the lamina propria into the epithelium. Free flow was assumed between all other internal as well as external boundaries.

### Combination of suction experiments and snap freezing

The tissue was further processed by removing fat and muscle tissue to trim it to ca. 2 – 3 cm thickness. It was then cut into pieces, placed in DMEM/F12 and prewarmed to 37 °C for 30 min before use. Suction experiments with the NIMBLE were performed with the same probes as described in the previous section (<u>suction measurement</u>). After reaching the maximal negative pressure, the probe together with the tissue samples were snap frozen in a liquid nitrogen chilled isopentane bath for ca. 1 min. Finally, the sample was embedded in an OCT and stored at - 20 °C until further processing within 2 weeks.

### Histochemical staining and imaging

The cryopreserved samples were sectioned with a cryostat (CryoStar NX50, Thermo Fisher Scientific) into 20 µm thin slices. During sectioning, the temperature was kept at -20 °C. Slides were thawed for 30 min at RT. Then, tissue sections were incubated for 5 min in freshly prepared 4% PFA in PBS and washed in PBS for 3 min. Thereafter, slides were permeabilized with 5% (v/v) Triton X100 containing 2% (w/v) BSA for 15 min and washed with a PBS solution at pH 7.4 for 3 min. Subsequently, the slides were incubated with the staining solution (1:1000 (v/v) iFluor^TM^ 488-conjugated phalloidin and 1:1000 (v/v) Hoechst 33342 in PBS containing 1% (w/v) BSA) for 30 min followed by a washing step in PBS for 3 min. In the last step, the slides were mounted with ProLong^TM^ Diamond Antifade Mountant and a coverslip.

### Imaging of stained samples

Fluorescent images of the sections were obtained using a Leica DMI 6000 B microscope with adaptive focus control (Leica Microsystems) at a magnification of 40X (HXC PL FL 40x/0.75, with CORR, numerical aperture: 0.4, dry, Leica Microsystems) using the LAS X software (Leica Microsystems) in the tile scan mode (manual and automatic focusing; range 100 µm) and were merged afterwards. Light was emitted by an EL6000 mercury metal halide bulb (Leica Microsystems). Images were detected by monochrome DFC365FX digital camera (12 bits, 1×1 BIN, Leica microsystems. Three channels per frame were used, i.e., Y5 for Cy5 (EX: 590 – 650 nm, DC: 660, EM: 662 – 738 nm), L5 for phalloidin conjugated iFluor^TM^ 488 (EX: 460 – 500 nm, DC: 505, EM: 512 – 542 nm, PN: 11504166), A4 for Hoechst 33342 (EX: 340 – 380 nm, DC: 400, EM: 450 – 490 nm).

### Evaluation of epithelial thickness

To evaluate the epithelial thickness, fluorescent images of the histochemically stained samples were used. The thickness was measured outside and in the center of the NIMBLE application site with the software ImageJ (National Institutes of Health) (Figure S1A). The distance was measured between the surface of the tissue and the basal membrane, recognizable by a dense layer of cells. In total 5 positions per region were measured on one sample.

### Evaluation of cell dimensions

For the evaluation of cell dimensions, fluorescent images of the histochemically stained samples were used. Single cells were randomly chosen in the mucosa region in the center (Figure S1B and D) and outside (Figure S1C and E) of the NIMBLE application side. The height and length of single cells were determined using the software ImageJ. In total 10 cells per region were measured on one sample.

### FEM epithelial thinning

The thinning of the epithelial was assessed in the FEM by extracting the Cauchy-Green tensor (stretch in the thickness direction) of the epithelium, 0.3 mm below the surface at the center of the specimen and at the outside of the specimen. The thickness was then calculated by multiplying the stretch, with a thickness of 1500 µm.

### FEM cell deformation

The cell deformation was extracted from the finite element simulation under the assumption of an affine deformation. The Cauchy-Green tensor was extracted for the corresponding elevation at the apex center in the lamina propria 0.3 mm under the surface. A geometric transformation was applied to the predefined cell outline using the local deformation gradient as the mapping function Eq. 10:

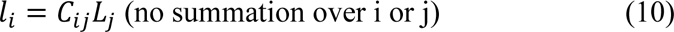

Therein *l_i_* represents the length in the deformed configuration, the length in the reference configuration and *C_ij_* the component of the Cauchy-Green tensor.

## Author contributions

DKC, DG and DS contributed equally to this work. All authors critically read and approved the paper.

## Data and materials availability

All data are available in the main text or the supplementary materials. Raw and source data are available in a repository database (doi: 10.5281/zenodo.14007674).

## Supplementary information

**Figure S1.**
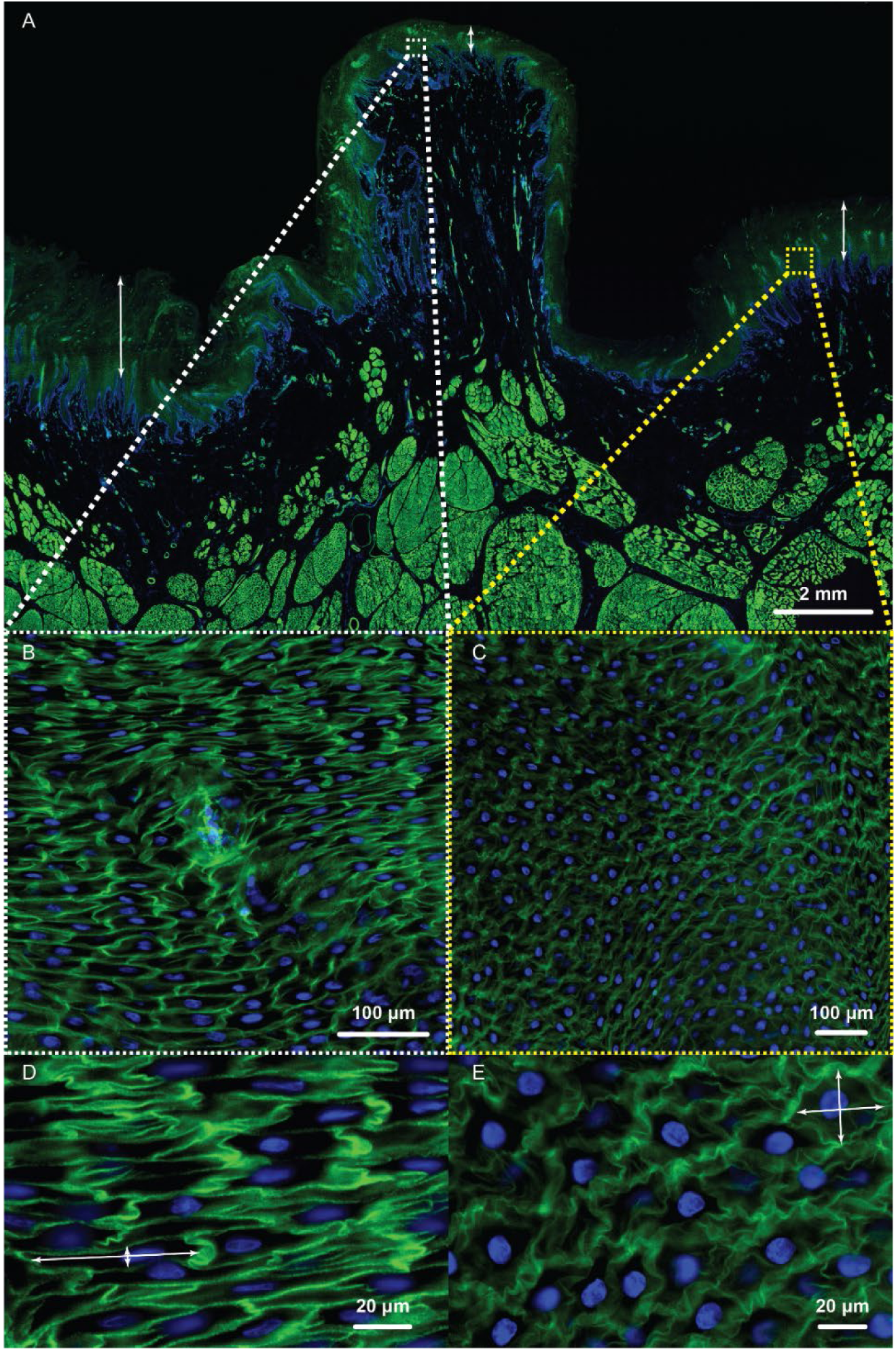
Example of histochemically stained porcine buccal tissue sample that was snap frozen during NIMBLE application. A series of magnifications were used for determining the epithelial thickness and cellular deformation. The cytoskeleton (F-actin, green) and nucleus (blue) were stained. A) An overview image and arrows indicate potential measurement points at the center of the NIMBLE application and an outside area that was assumed not stretched as reference. B) and C) the magnification at the center and outside of the application side, respectively. D) The magnification of the center with elongated cells. The white arrows show examples of measurement points for cell deformation measurements. E) Further magnification of C) showcases the circular cell structure outside the application side.

**Figure S2.**
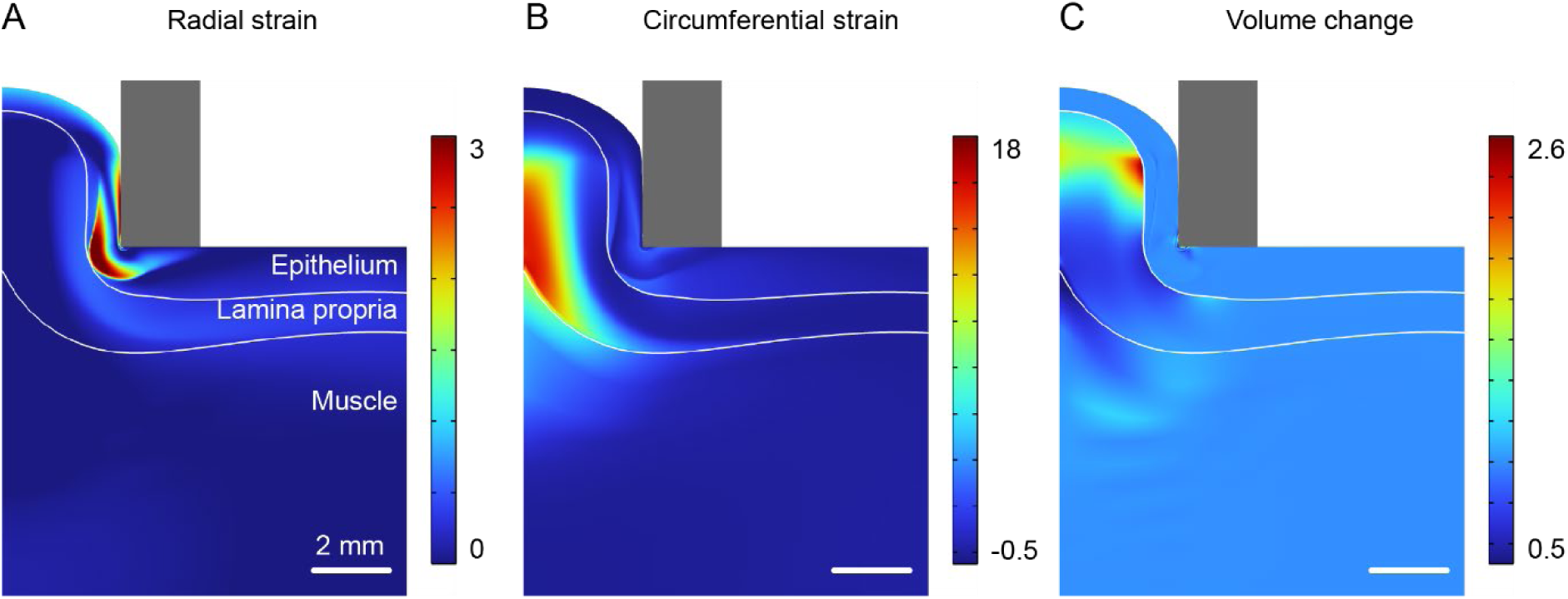
Deformation behavior predicted by the finite element simulations for a deformation height of 4 mm. A) Strains in the radial direction and B) in the circumferential direction and C) corresponding volume change. Note that no volume change was predicted in the epithelium due to the assumption of incompressibility. Scale bars = 2 mm.

**Figure S3.**
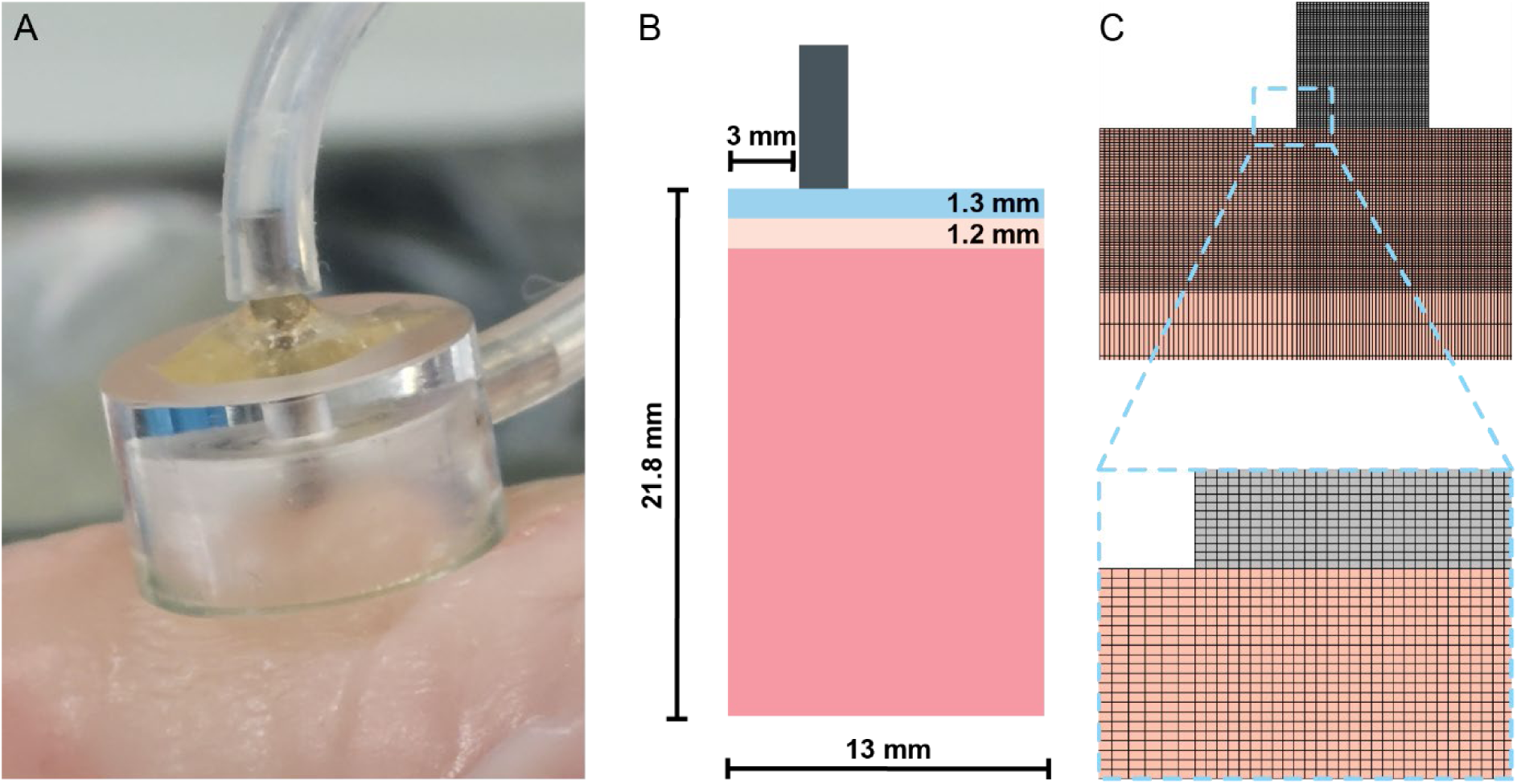
Experimental application and FEM architecture. A) NIMBLE probe (3 mm radius) applied on porcine buccal mucosa at the required pressure. B) Layer structure and dimensions used in the FEM model of the epithelium (blue), lamina propria (orange), and muscle (red) as well as the wall of the NIMBLE specimen (gray). C) Mesh of the NIMBLE probe (gray) and the tissue (orange). The magnification highlights the fine mesh used at the interface of both structures.

**Table S1.**
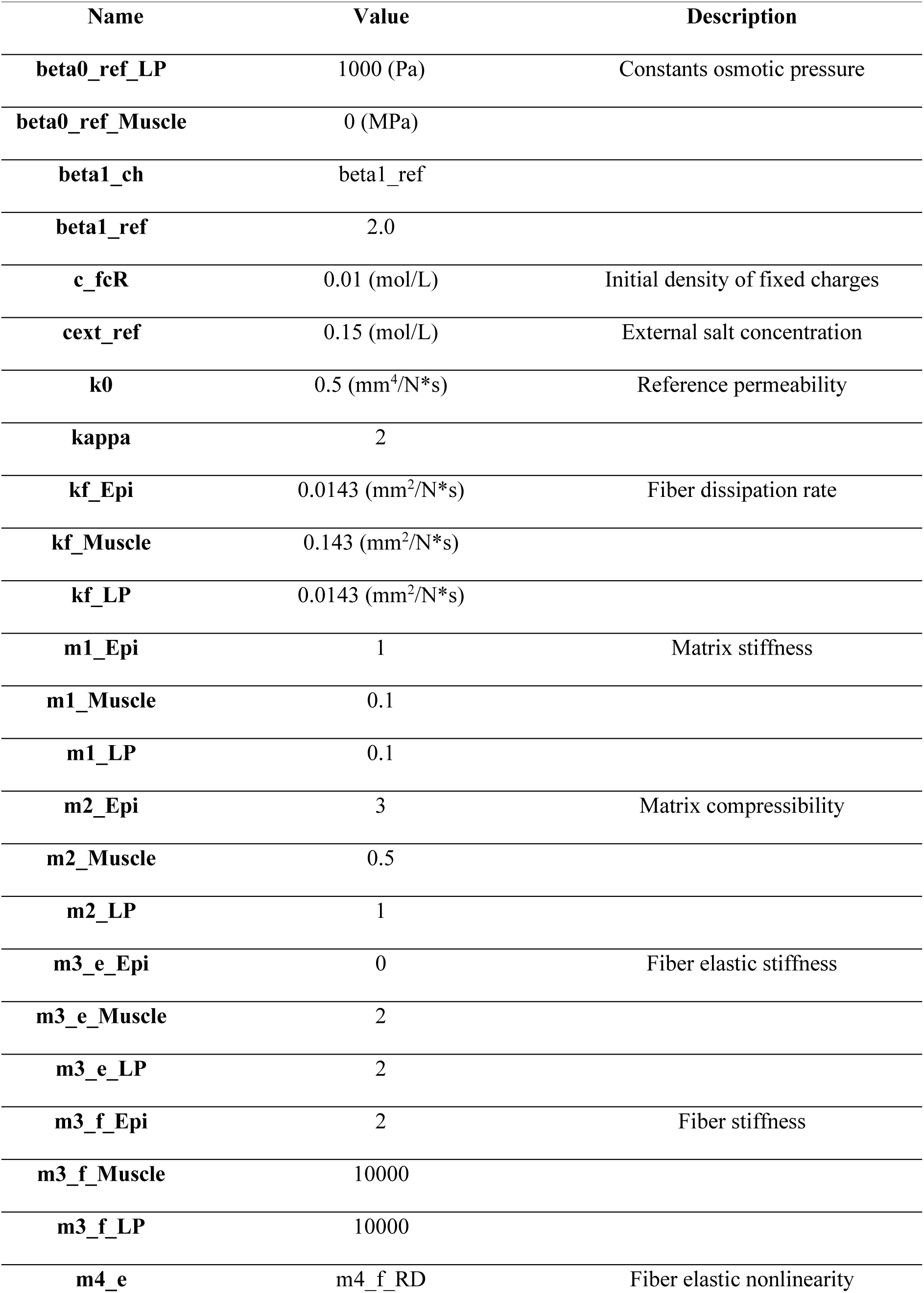
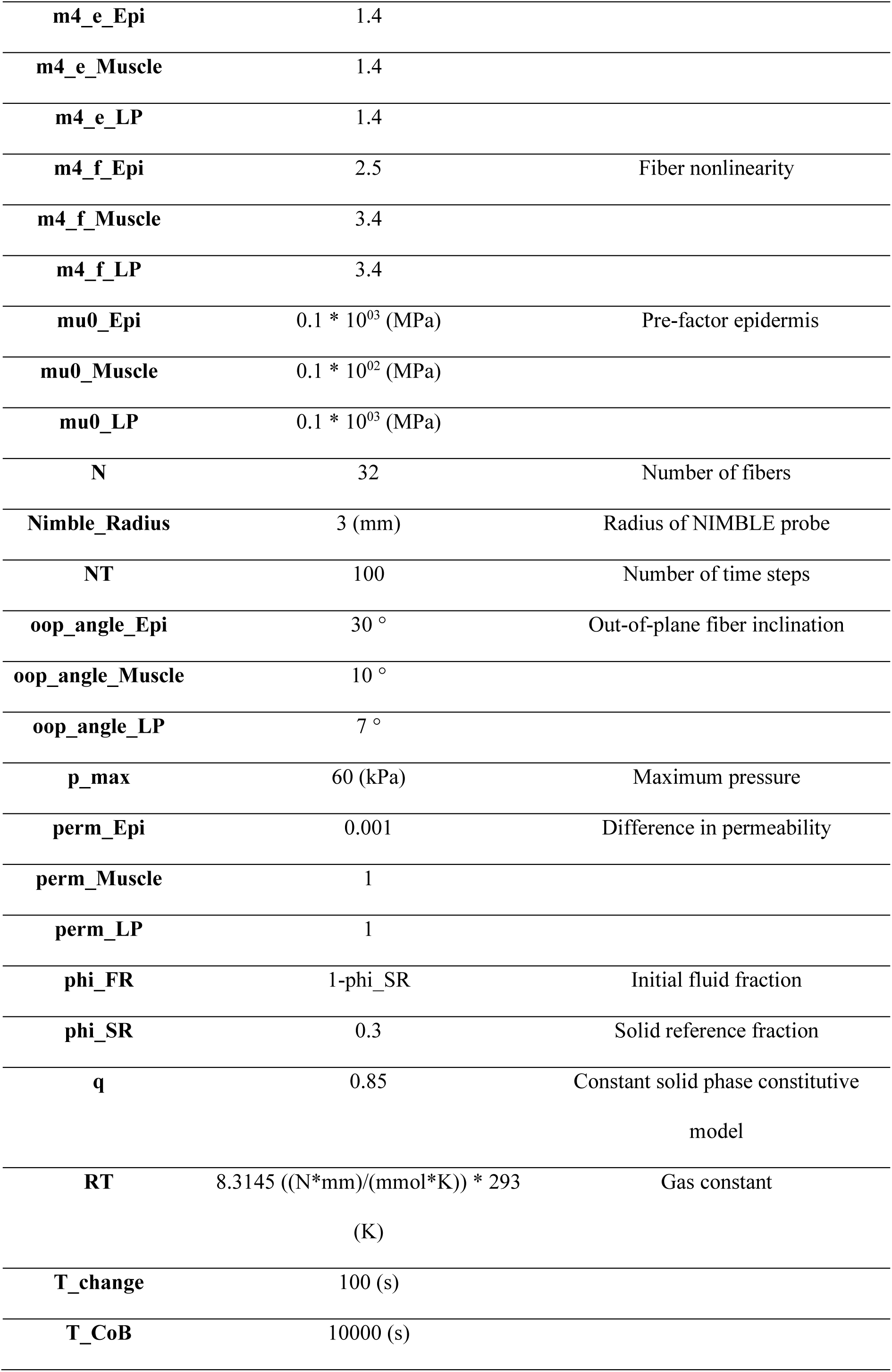
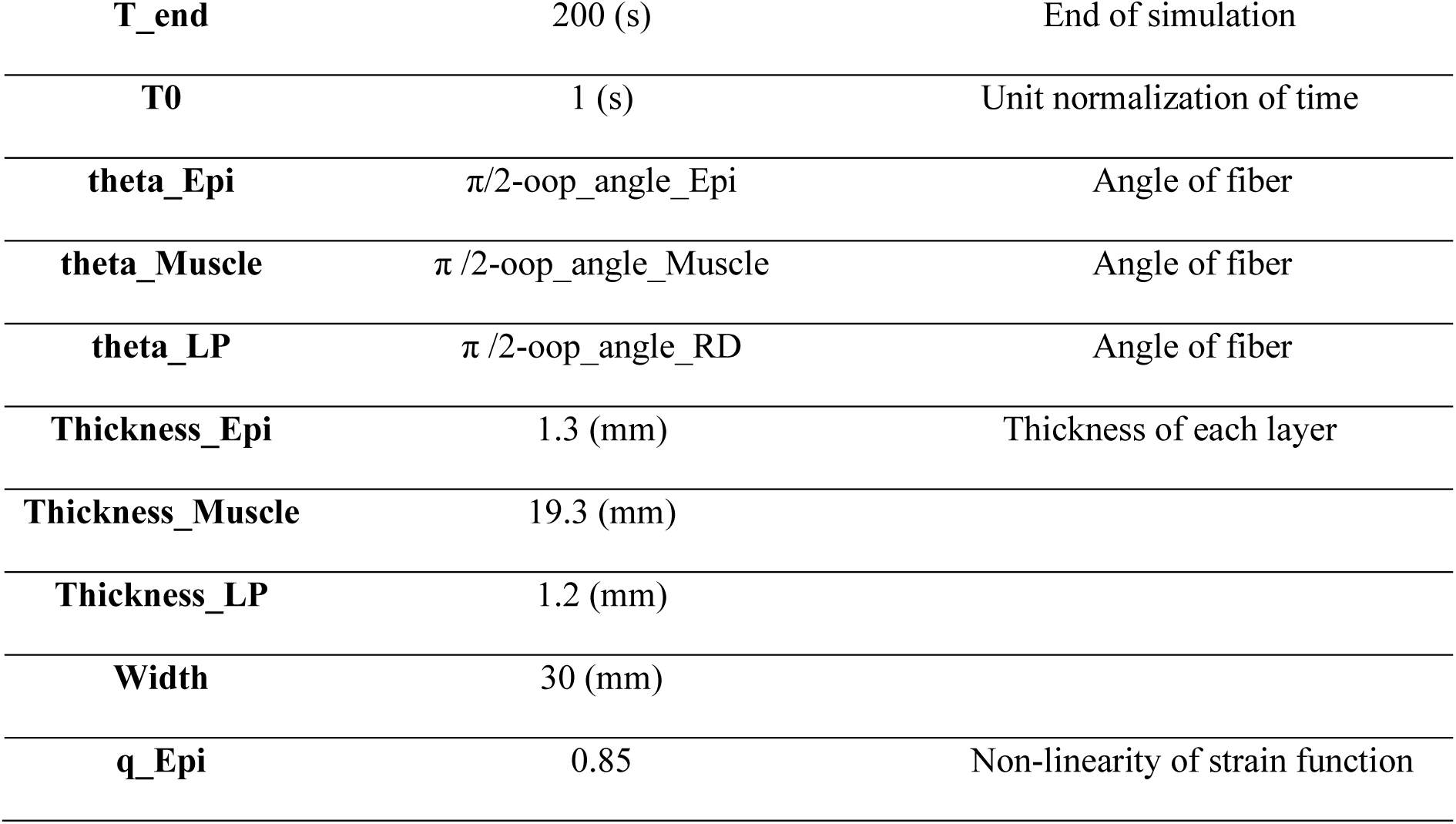
Optimized parameters of the trained FEM.

**Table S2.** Young’s moduli obtained in computational uniaxial test at high and low strain of individual tissue layers.

| **Young's modulus (kPa)** |  |  |
| --- | --- | --- |
| **Strain** | Low (1 – 10%) | High (40 – 50%) |
| **Epithelium** | 0.09 | 0.16 |
| **Lamina Propria** | 0.09 | 45.1 |
| **Muscle** | 0.89 | 4510 |

